# Mitochondrial dsRNAs activate PKR and TLR3 to promote chondrocyte degeneration in osteoarthritis

**DOI:** 10.1101/2020.06.17.156323

**Authors:** Sujin Kim, Keonyong Lee, Yong Seok Choi, Jayoung Ku, Yun Jong Lee, Yoosik Kim

## Abstract

Protein kinase R (PKR) is an immune response protein that becomes activated by long double-stranded RNAs (dsRNAs). Several studies reported the misactivation of PKR in patients of degenerative diseases including primary osteoarthritis (OA). However, the molecular identity of PKR-activating dsRNAs remains unknown. Here, we investigate the role of mitochondrial dsRNAs (mt-dsRNAs) in the development of OA. We find that in response to OA-mimicking stressors, cytosolic efflux of mt-dsRNAs is increased, leading to PKR activation and subsequent induction of inflammatory cytokines and apoptosis. Moreover, mt-dsRNAs are exported to the extracellular space where they activate toll-like receptor 3. Elevated expression of mt-dsRNAs in the synovial fluids of OA patients further supports our data. Lastly, we show that autophagy protects chondrocytes from mitochondrial dysfunction partly by removing cytosolic mt-dsRNAs. Together, these findings establish the PKR-mt-dsRNA as a critical regulatory axis in OA development and suggest mt-dsRNAs as a potential target in fighting OA.

## INTRODUCTION

Protein kinase R (PKR) is an omnipresent enzyme expressed in vertebrates. It is a member of the innate immune response system that recognizes long double-stranded RNAs (dsRNAs) from viruses^1,2^. In this context, PKR binds to dsRNAs longer than 33 base-pairs (bp), which leads to dimerization and subsequent autophosphorylation of the enzyme^3–5^. Phosphorylated PKR (pPKR) becomes an active kinase and phosphorylates a range of substrates such as p53, STAT, and eIF2α to mediate stress response and suppress global translation^6–8^. Moreover, PKR also activates multiple downstream signaling pathways such as JNK, MAPK, and NF-κB to initiate a cellular response to various external stimuli^7^. Recently emerging evidence has established that PKR is a multifunctional protein whose activation status needs to be closely monitored to maintain proper cell physiology. For example, PKR activation is regulated in a cell cycle-dependent manner where its phosphorylation is increased during the M-phase to control global translation and orchestrate mitotic processes^9,10^. In addition, PKR is involved in healthy brain function as knockdown of PKR resulted in improved cognitive function in mice^11,12^. Overactivation of PKR is a hallmark of numerous degenerative diseases ranging from Parkinson’s disease, Alzheimer’s disease, and even primary osteoarthritis (OA)^12–17^. PKR activation during apoptosis further supports the idea that PKR may play a key regulatory function in the onset and progression of degenerative diseases.

Primary OA is a critical degenerative disorder of the diarthrodial synovial joints that affects a considerable part of the adult population. It is the most common type of arthritis and occurs mostly on the knee as well as hip and hands^18^. Previous studies have shown that increased levels of metalloproteases (MMPs) such as MMP-1 and MMP-13 as well as disintegrins and metalloproteinases are partly responsible for the degeneration of collagens and proteoglycans^19^. Even though OA has been considered a non-inflammatory arthropathy, numerous proinflammatory cytokines play a role in the pathogenesis where they may drive the induction of MMPs and ADAMs^20^. Recently, studies have shown that pattern recognition receptors (PRRs) such as toll-like receptor 3 (TLR3) and PKR are involved as upstream regulators that induce the expression of cytokines^21,22^. Indeed, the increased level of pPKR was reported in damaged chondrocytes, and dsRNAs released from damaged chondrocytes induced TLR3 activation to promote cartilage degeneration^22^. Yet, the molecular identity of these dsRNAs that can activate TLR3 and PKR remains unknown.

Long dsRNAs are a signature of viral infection as these RNAs are generated as byproducts of viral replication and through bidirectional transcription of viral DNAs^23^. They are abundantly present in DNA and positive-sense single-stranded RNA viruses^24^. When expressed in mammalian cells, these dsRNAs are recognized by PRRs, which leads to suppression of global translation, induction of interferons and proinflammatory cytokines, and eventually, cell death ^25^. However, increasing evidence suggests that human cells naturally encode cellular dsRNAs that can regulate antiviral machinery. Previous studies have shown that short-interspersed nuclear elements (SINEs), mostly Alus in primates, can be transcribed and generate dsRNAs^26,27^. As Alu elements constitute over 10% of the human genome, multiple Alus are embedded in the 3′ UTRs and intronic regions of most protein-coding genes and are co-transcribed together with the host gene^28–30^. When transcribed, these Alu elements can bind to each other and form an extended double-stranded secondary structure, which is recognized by PKR, leading to PKR activation under physiological conditions^31,32^. In addition, numerous high-throughput studies have shown that other retrotransposons such as long-interspersed nuclear elements (LINEs) and endogenous retroviral elements (ERVs) can also be transcribed into long dsRNAs that can activate RNA sensors of the innate immune response proteins such as PKR and melanoma-differentiation associated protein 5 (MDA5)^32–34^. More importantly, misactivation of PRRs and endogenous dsRNAs are closely associated with numerous degenerative diseases^35^. It has been shown that decreased expression of RNase III enzyme Dicer can promote age-related macular degeneration through the accumulation of Alu transcripts^36^. In addition, inducing LINE-1 expression by altering the methylation status of its promoter can contribute to OA development^37^.

A recent study employed formaldehyde cross-linking and immunoprecipitation followed by high-throughput sequencing (fCLIP-seq) to identify PKR-interacting cellular dsRNAs in human cells^32^. In addition to retrotransposons, a major contributor of the RNA interactor of PKR is mitochondrial double-stranded RNAs (mt-dsRNAs). These mt-dsRNAs are generated due to the bidirectional transcription of the circular mitochondrial genome, which results in the transcription of long complementary RNAs^38^. Moreover, mt-dsRNAs can also activate MDA5 in mice and can be secreted in exosomes to activate TLR3 in liver cells during alcoholic stress^39,40^. Considering that the mitochondrial dysfunction is frequently reported in OA patients and that mt-dsRNAs can activate multiple PRRs to induce expression of inflammatory cytokines and apoptosis^32,39,41–43^, the role of mt-dsRNAs during the onset and progression of OA needs further investigation.

In this study, we set out to examine whether mt-dsRNAs are responsible for PKR and TLR3 activation to promote chondrocyte degeneration during the pathogenesis of primary OA. Our results show that in response to mitochondrial stress, mt-dsRNAs strongly interact with PKR and activate the PKR signaling pathway. More importantly, downregulation of PKR or mt-dsRNAs can attenuate the expression of key MMPs and cytokines that are associated with OA development. In addition, mt-dsRNA-PKR pathway is not limited to mitochondrial stress as the cytosolic efflux of mt-dsRNAs, and subsequent PKR activation also occur during the senescence of chondrocytes, a key process in the early stages of OA. Moreover, we find that mt-dsRNAs can be exported outside the cells, where they act as intercellular signaling cues to activate TLR3 signaling in the neighboring cells. Lastly, we show that autophagy alleviates OA conditions partly by removing cytosolic mt-dsRNAs. Collectively, our findings establish mt-dsRNAs as the unknown cellular dsRNAs released from damaged chondrocytes, uncover the key regulatory function of mt-dsRNAs during the onset and progression of OA, and even suggest a potential target to fight OA.

## Results

### Inhibition of mitochondrial respiratory chain activates PKR

Mitochondrial dysfunction has been associated with primary OA development as it results in increased production of reactive oxygen species (ROS), which subsequently leads to inflammation and decreased ability of chondrocyte synthesis. We sought out to investigate how damaged mitochondria could result in the activation of inflammatory signaling and contribute to OA development. In particular, we asked whether mitochondrial dysfunction could result in the activation of PKR, which can disrupt cell homeostasis by suppressing translation and initiating apoptotic programs. To trigger mitochondrial dysfunction, we treated SW1353 chondrosarcoma cells with oligomycin A (oligoA) to inhibit ATP synthase of the mitochondrial respiratory chain complexes (MRCs), as described in previous studies^44,45^. Using JC-1 dye, we confirmed that oligoA treatment damages mitochondria, which is reflected in the disruption of mitochondrial membrane potential (Fig. 1a). Consistent with previous studies, we found that oligoA treatment resulted in significant cell death accompanied by induction of senescence-associated secretory phenotype (SASP) factors as well as interferons (IFNs) and interferon-stimulated genes (ISGs) (Fig. 1b, c). Using this system, we asked whether PKR plays a role in the cellular response to mitochondrial dysfunction. We examined the activation of PKR through immunocytochemistry and found that oligoA treatment resulted in increased phosphorylation of PKR and its downstream factor eIF2α, indicating that disruption of MRCs results in activation of the PKR signaling pathway (Fig. 1d). To rule out the possibility that the observed phenomena are specific to cancer cells, we verified our results using immortalized human chondrocyte cell line CHON-001. OligoA treatment resulted in cell death with increased expression of SASP factors, IFNs, and ISGs in CHON-001 cells as well (Supplementary Fig. 1a, b). Moreover, PKR and its downstream signaling are also activated by oligoA in CHON-001 cells (Supplementary Fig. 1c).

**Fig. 1.**
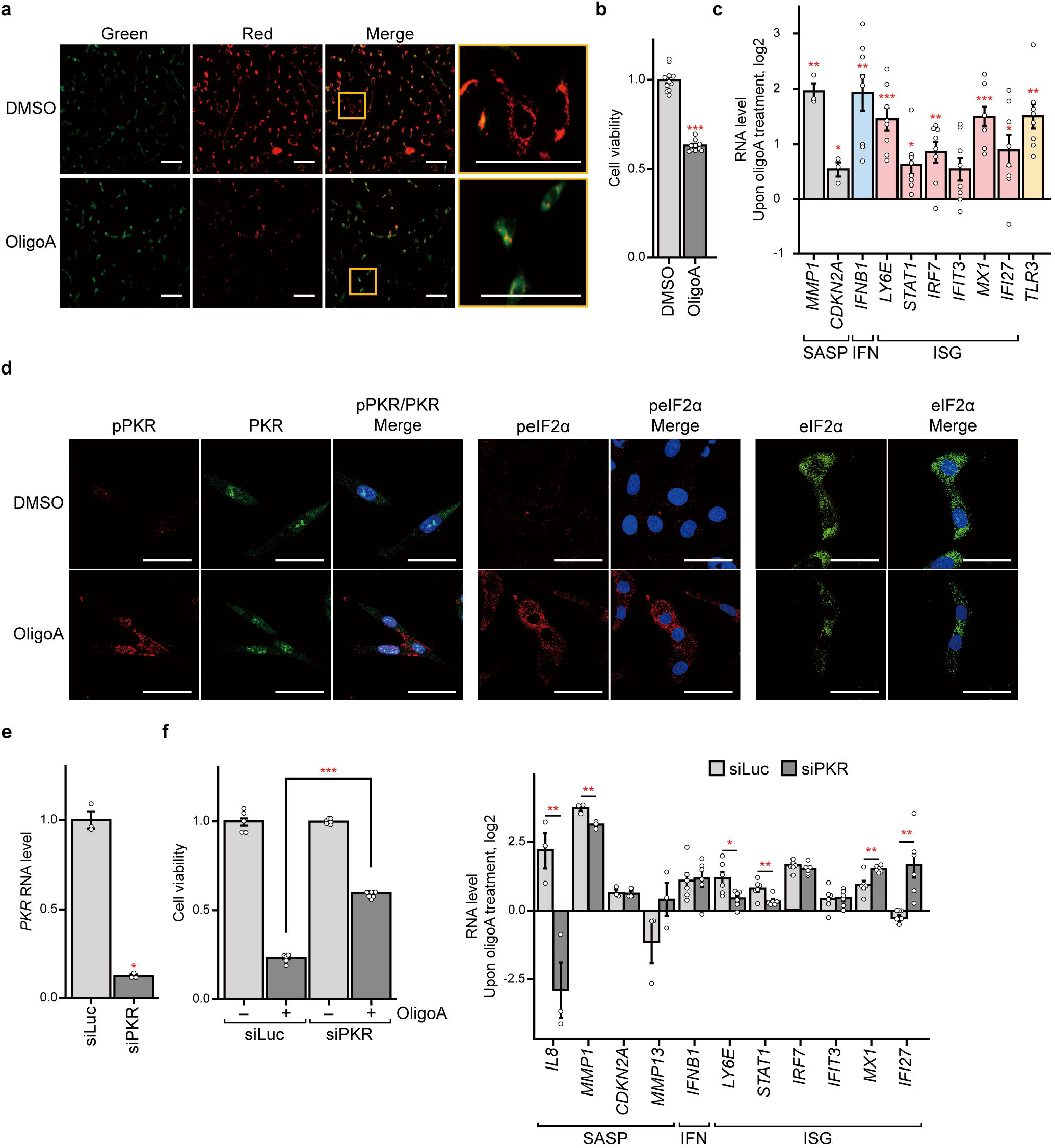
MRC inhibition activates the PKR signaling pathway to promote chondrocyte degeneration. **a-c** Inhibition of MRC by oligoA results in disruption of mitochondrial membrane potential, assessed **a** by the increased green fluorescent signal from the JC-1 dye, **b** decreased cell viability (*n* ≥ 10), and **c** induction of SASP factors, IFN, and ISGs (*n* = 3 for SASP factors and *n* = 8 for IFN and ISGs). Scale bar is 100 μm in **a**. **d** OligoA treatment results in the activation of PKR and phosphorylation of its downstream substrate eIF2α. Scale bar is 50 μm. **e-g** Using RNA interference, **e** PKR expression can be knocked down (*n* = 3), which partially rescues the effect of oligoA treatment on **f** cell viability (*n* ≥ 5) and **g** induction of SASPs and ISGs (*n* = 3).

To establish that PKR plays a key role in response to oligoA treatment, we used RNA interference to knockdown PKR expression and examined its effect (Fig. 1e). In PKR-deficient cells, cell death from oligoA treatment was rescued substantially (Fig. 1f). Moreover, RNA levels of key OA-associated factors such as *IL8, MMP1, LY6E,* and *STAT1* were significantly decreased when PKR was suppressed (Fig. 1g). This indicates that PKR is a critical regulator in the disruption of cell homeostasis associated with the inhibition of MRC.

### MRC inhibition induces cytosolic efflux of mt-dsRNAs to activate PKR

We next sought out to identify the activating cue responsible for PKR phosphorylation during MRC inhibition. A recent study has shown that decreased expression of PNPase resulted in the increased cytosolic efflux of mt-dsRNAs, which are recognized by MDA5, leading to MDA5 oligomerization and IFN induction^39^. We examined the expression of PNPase and hSUV3 upon oligoA treatment, but their mRNA expressions did not change (data not shown). In addition, the level of mtRNAs was not affected by oligoA. Considering that oligoA alters the mitochondrial membrane potential, we hypothesize that it may induce cytosolic efflux of mtRNAs, which may account for the increased phosphorylation of PKR. To test, we examined PKR-mtRNA interaction via fCLIP approach and found that PKR strongly interacts with mtRNAs when oligoA was treated (Fig. 2a). Although the expression of total mtRNA remained unchanged, the level of cytosolic mtRNAs was significantly increased, which could account for the observed mtRNA-PKR interactions (Fig. 2b). To measure the cytosolic mtRNA expression, we first isolated the free cytosolic compartment from organelles (membrane fraction) and the nucleus and extracted RNAs on the three compartments. Successful fractionation was confirmed by examining marker protein patterns via western blotting (Supplementary Fig. 2a). We further performed strand-specific reverse transcription to examine the mtRNAs that originated from heavy and light strands. We found that both heavy and light mtRNAs levels in the cytosol were increased by oligoA treatment (Fig. 2c), suggesting the potential dsRNA generation through the interaction of these complementary RNAs. We also confirmed that oligoA treatment resulted in increased cytosolic mtRNA expression in CHON-001 cells as well (Supplementary Fig. 2a, b). Overall, these results suggest that MRC inhibition via oligoA results in the cytosolic efflux of mt-dsRNAs, where they are recognized by PKR, leading to PKR autophosphorylation (Fig. 2d).

**Fig. 2.**
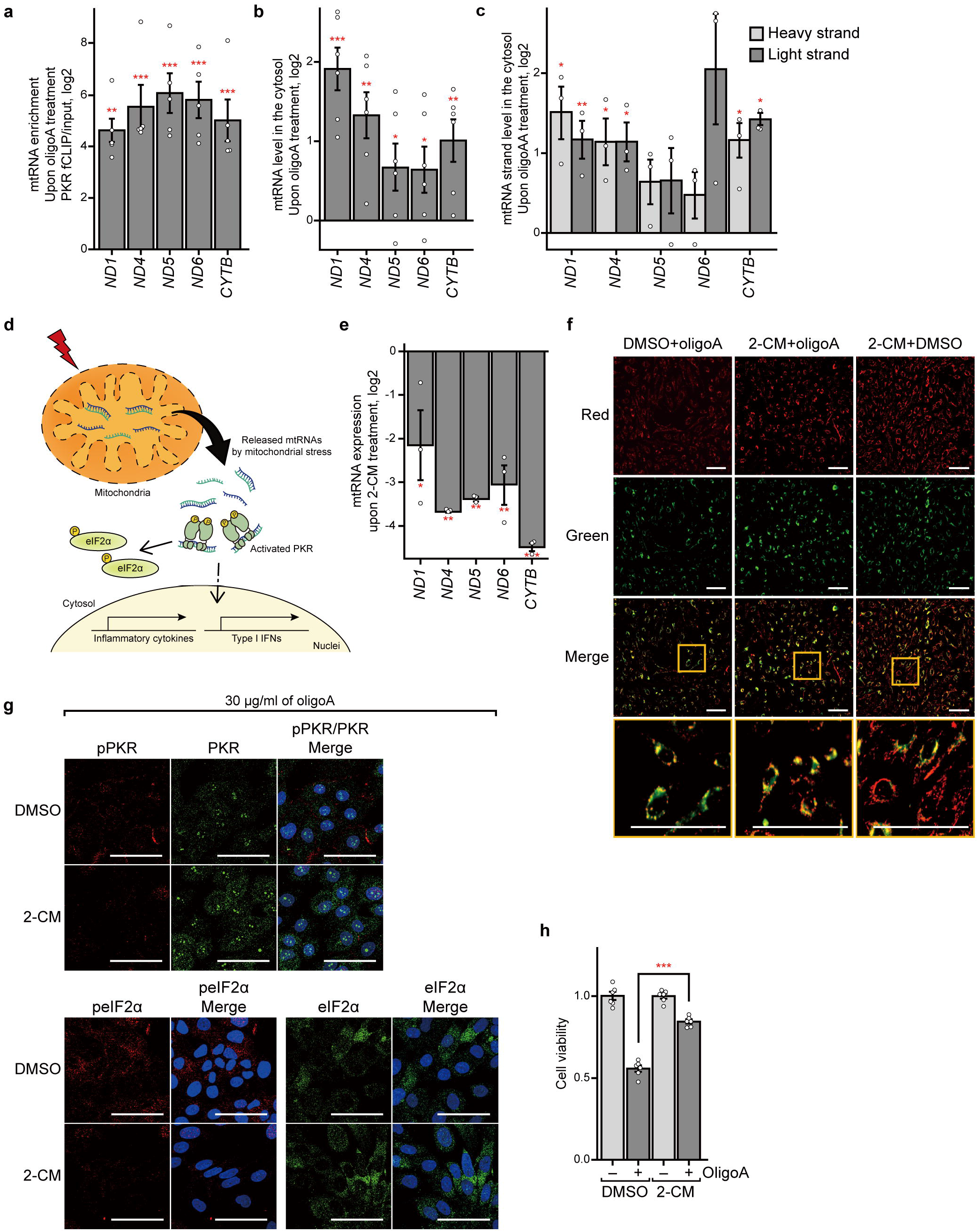
Mitochondrial RNAs interact with PKR during MRC inhibition. **a** PKR fCLIP-qPCR analysis reveals strong interaction between most mtRNAs and PKR upon oligoA treatment. An average of 5 biological replicates is shown with error bars indicating s.e.m. **b, c** Increased PKR-mtRNA interaction is likely due to cytosolic efflux of mtRNAs. **b** Total and **c** strand-specific mtRNA expressions in the free cytosolic fraction is measured by qRT-PCR. *n* = 6 for total mtRNA and *n* = 3 for strand-specific mtRNA expressions and error bars denote s.e.m. **d** A schematic of the model where cytosolic mtRNAs activate PKR during mitochondrial stress. **e** mtRNA expression is suppressed by treating cells with 2-CM. qRT-PCR on total mtRNAs is shown with *n* = 3 and error bars denote s.e.m. **f** OligoA treatment still disrupts the mitochondrial membrane potential even in 2-CM pretreated cells. Scale bar, 100 μm. **g, h** In cells with decreased expression of mtRNAs, MRC inhibition does not activate **g** the PKR signaling pathway and show increased **h** cell viability (*n* = 6). Scale bar is 50 μm in **g**.

In addition to mt-dsRNAs, human cells encode various types of non-coding RNAs and repeat RNAs that can activate PKR. We examined whether mt-dsRNAs are a key contributor in activating PKR during MRC inhibition. We first pretreated cells with 2′-C-Methyladenosine (2-CM), a small molecule inhibitor of mitochondrial RNA polymerase, to decrease the expression of mtRNAs. We found that 2-CM treatment in SW1353 cells decreased mtRNA expression by 80~90% (Fig. 2e). We confirmed that 2-CM treatment alone does not disrupt the mitochondrial membrane potential (Fig. 2f). The disruption of mitochondrial membrane potential by oligoA was similar, regardless of 2-CM treatment (Fig. 2f). Under this condition, we applied oligoA and examined its effect on PKR signaling. Interestingly, we found that oligoA treatment failed to activate PKR in 2-CM pretreated cells (Fig. 2g). In addition, eIF2α phosphorylation was reduced significantly in cells with decreased expression of mtRNAs (Fig. 2g). More importantly, 2-CM pretreatment resulted in significant rescue in cell viability (Fig. 2h), similar to that of PKR-deficient cells. Therefore, our data clearly suggest that mtRNAs are the key source of PKR activation during MRC inhibition.

### Senescence-eliciting stressors also induce PKR activation via mt-dsRNAs

Chondrocyte senescence is a critical event contributing to the early development of OA by disrupting the balance between matrix anabolism and catabolism^46,47^. A recent study has shown that induction of senescence by prolonged exposure to doxorubicin (Dox) or hydrogen peroxide (H_2_O_2_) resulted in increased microRNA-204 expression and subsequent cessation of the anabolic pathway^48^. We asked whether PKR and mt-dsRNAs also contribute to the chondrocyte senescence. SW1353 cells were exposed to Dox or H_2_O_2_ for three or four days, respectively, and senescence was confirmed using senescence-associated β-galactosidase (SA-β-gal) staining (Fig. 3a). Consistent with previous studies, we confirmed that chondrocyte senescence induced key OA-associated factors such as *IL8* and *MMP1* in addition to other SASP markers (Fig. 3b). Interestingly, we found that senescent cells showed strong pPKR and peIF2α signals (Fig. 3c, d, and Supplementary Fig. 3a). Particularly, extended exposure to H_2_O_2_ resulted in more pronounced activation of the PKR signaling pathway.

**Fig. 3.**
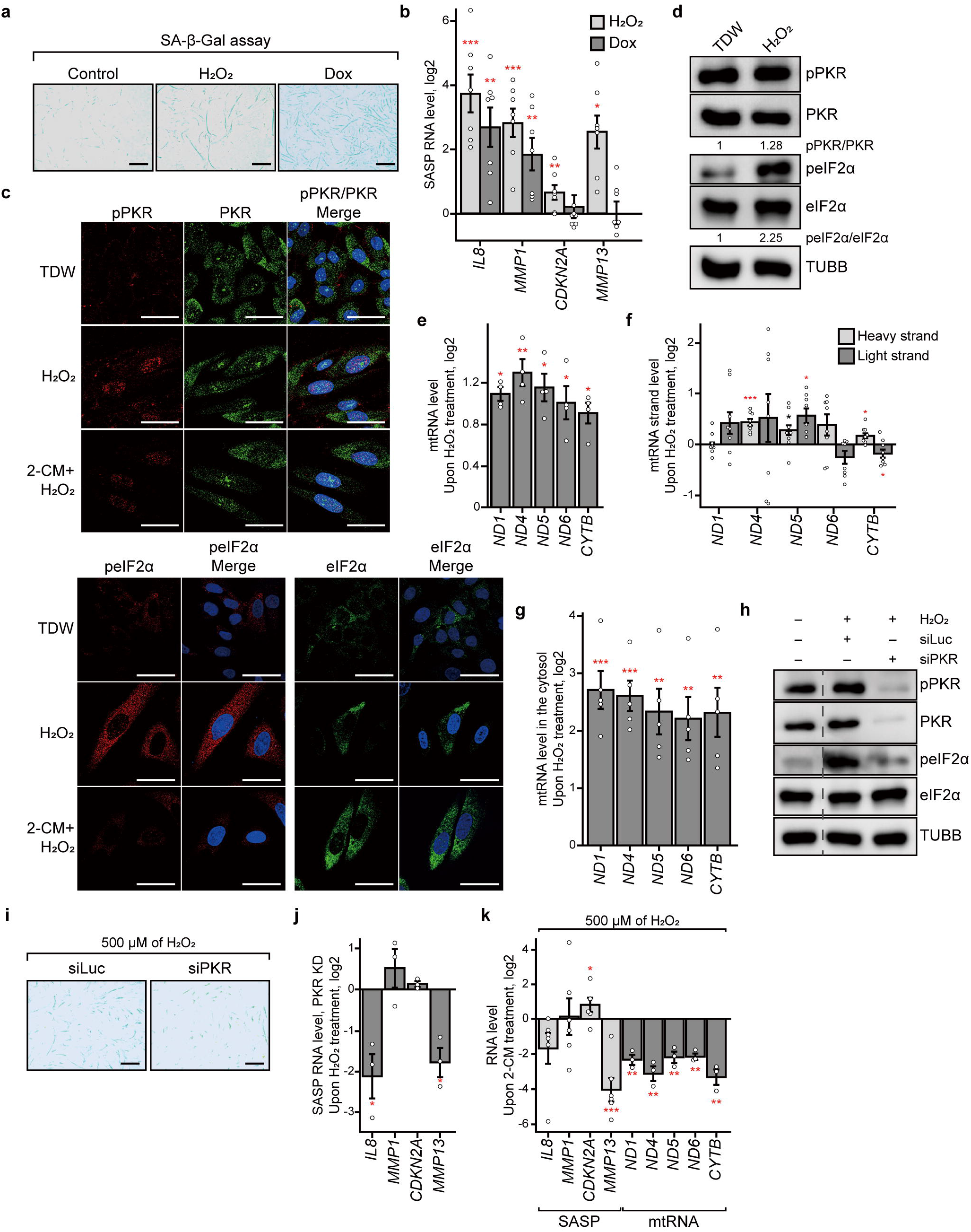
Chondrocyte senescence eliciting stress conditions also activate the PKR signaling pathway via mtRNAs. **a** SA-β-Gal staining indicates successful senescence induction by H_2_O_2_ and Dox treatment. Scale bar, 500 μm. **b** Relative mRNA expressions of SASP factors in response to H_2_O_2_ and Dox treatment. Average of 7 is shown with error bars representing s.e.m. **c, d** Activation of PKR and phosphorylation of eIF2α during senescence are confirmed using **c** immunocytochemistry and **d** western blotting. Scale bar, 50 μm. **e, f e** Total (*n* = 4) and **f** strand-specific mtRNA (*n* = 8) expression upon prolonged exposure to H_2_O_2_. Note the pronounced increase in the antisense transcripts. **g** The expression of free cytosolic mtRNAs is also increased by H_2_O_2_ treatment. *n* = 5 and error bars denote s.e.m. **h-j** In PKR-deficient cells, H_2_O_2_ treatment results in decreased **h** eIF2α phosphorylation, **i** SA-β-Gal activity, and **j** *IL8* and *MMP13* mRNA levels (*n* = 3). In **i**, the scale bar denotes 500 μm. **k** Similarly, in mtRNA-deficient cells, H_2_O_2_ treatment results in decreased *MMP13* mRNA expressions (*n* = 6).

We further investigated whether mt-dsRNAs are also responsible for PKR activation in promoting senescence. We found that prolonged exposure to H_2_O_2_ increased the level of mtRNAs (Fig. 3e). Using strand-specific RT-qPCR, we further examined this change and found that for most mitochondrial genes tested, a more significant increase in antisense mtRNAs (light-strand RNAs for the most part and heavy-strand RNA for the ND6 gene) was observed (Fig. 3f). Considering that the protein-coding sense mtRNAs are more abundant than that of the counterpart antisense RNAs, this increase in antisense mtRNAs would likely result in increased mt-dsRNA expression. We further performed subcellular fractionation and found that cytosolic mtRNA levels were significantly increased upon H_2_O_2_ treatment, which could account for the increased PKR phosphorylation (Fig. 3g). Indeed, reduction of mtRNA expression by 2-CM pretreatment abrogated pPKR and peIF2α signals while it did not affect total PKR and eIF2α expression (Fig. 3c). Of note, we confirmed similar but weaker effects when Dox was used to induce senescence. One possibility is that for Dox, expression of nc886, a non-coding RNA that inhibits PKR activation, may have affected PKR activation, as shown in a previous study^49^. Regardless, the main finding that cellular senescence resulted in increased total and cytosolic levels of mtRNAs is still valid (Supplementary Fig. 3a-d).

To establish PKR as a key senescence-associated factor for OA development, we knocked down PKR expression and analyzed its effect on senescence phenotypes. We found that in PKR-deficient cells, induction of peIF2α by prolonged exposure to H_2_O_2_ was significantly reduced (Fig. 3h). More importantly, SA-β-gal activity and *IL8* and *MMP13* expressions were decreased in PKR knockdown cells (Fig. 3i, j, and Supplementary Fig. 3e). Interestingly, a senescence marker *CDKN2A* mRNA level was not affected, suggesting that PKR has more pronounced effects on OA-associated factors. In addition, we analyzed the impact of decreasing mtRNA expression on senescence phenotypes. 2-CM pretreatment also resulted in a significant decrease in *MMP13* mRNA expression and, although statistically insignificant, *IL8* and *MMP1* mRNA, indicating that mt-dsRNAs are a key activator of PKR to promote OA development during chondrocyte senescence (Fig. 3k).

### mt-dsRNAs act as intercellular signaling cue by activating TLR3

One explainable factor of OA progression is dsRNAs released from damaged chondrocytes that promote cartilage degeneration by activating TLR3 signaling pathway^22^. However, the molecular identity of TLR3-activating dsRNAs in the synovial fluid remains unknown. We asked whether mt-dsRNAs can be released outside the cell to the synovial fluid and activate TLR3 in the neighboring cells during OA development (Fig. 4a). To test, we induced mitochondrial dysfunction using oligoA and collected and extracted RNAs in the media after 24 h. Compared to control cells with DMSO treatment, oligoA treated cells showed significantly higher levels of mtRNAs in the media (Fig. 4b and Supplementary Fig. 4a). Of note, extracellular mtRNA level was normalized to that of *GAPDH* mRNA in order to account for variable RNA expression and possible cell contaminants. To confirm that these extracellular mtRNAs can activate TLR3, we treated oligoA in TLR3-deficient cells and examined the expression of TLR3 downstream genes. We found that knockdown of TLR3 significantly decreased *IRF3* as well as several ISGs such as *LY6E, IRF7,* and *IFI27* (Fig. 4c). Of note, as shown earlier and by previous studies, mt-dsRNAs can induce interferons and ISGs via PKR and MDA5. Therefore, we cannot pinpoint the ISGs that are regulated by the TLR3-mt-dsRNA pathway alone. Nevertheless, our data indicate that TLR3, which recognizes dsRNAs in the extracellular space, also contributes to the ISG induction by oligoA treatment.

**Fig. 4.**
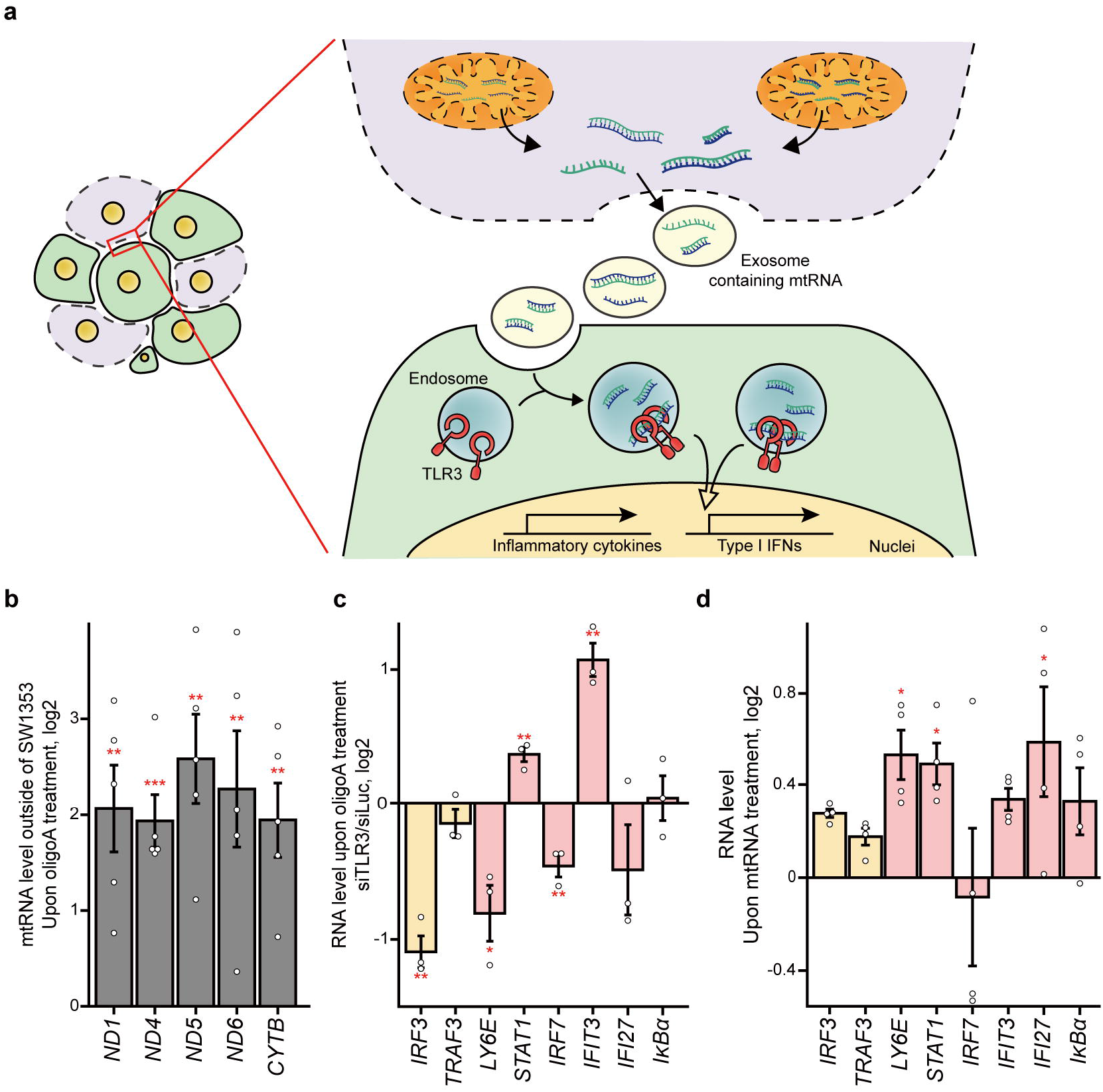
mtRNAs can act as autocrine-like signaling cues to activate TLR3. **a** A model for the export of mt-dsRNAs through exosomes, which can activate TLR3 in the neighboring cells to promote cartilage degeneration. **b** The expression of mtRNAs in media is significantly increased upon MRC inhibition. *n* = 5 and error bars denote s.e.m. **c** Induction of TLR3 signaling targets and several key ISGs is significantly decreased in TLR3-deficient cells. *n* = 3 and error bars denote s.e.m. **d** TLR3 signaling and ISGs can be induced by treating chondrocytes with RNAs extracted from isolated mitochondria. *n* = 4 and error bars denote s.e.m.

To rule out the possibility that the observed effects are due to direct activation of TLR3 by oligoA, we treated cells with mtRNAs directly and examined the downstream ISG induction. To collect mtRNAs, we isolated mitochondria from SW1353 cells, treated them with RNase A to remove any RNAs on the mitochondrial surface, and then extracted RNAs. We found that treating cells with RNAs isolated from mitochondria had similar effects on the induction of TLR3 downstream factors and ISGs (Fig. 4d). Therefore, upon mitochondrial stress, mt-dsRNAs are released to the extracellular space where they can act as autocrine-type signaling molecules to activate TLR3 in the neighboring chondrocytes and may induce local inflammation.

### Cellular efflux of mtRNAs is increased in OA patients

The importance of mt-dsRNAs in the activation of PKR and subsequent induction of SASP factors as well as several ISGs related to OA development led us to examine cartilages and primary chondrocytes from the patients. We acquired pieces of cartilages from seven OA patients undergoing total knee replacement. The mean age of the patients was 74.3 ± 5.01 years. We minced articular cartilage tissues into small pieces using surgical blades for tissue culture (Fig. 5a). We then induced mitochondrial stress by treating the cartilages with oligoA and analyzed the extracellular efflux of mt-dsRNAs. Consistent with the cell line experiments, we detected an increased expression of most mtRNAs in the media from the stressed cartilages. The expressions were quite variable from sample to sample, but we consistently observed increased efflux of mtRNAs into the media (Fig. 5b). This suggests that the damaged/stressed articular chondrocytes releases mtRNAs into the extracellular space where they may act as intercellular dsRNAs to trigger TLR3 to promote cartilage degeneration as suggested previously.

**Fig. 5.**
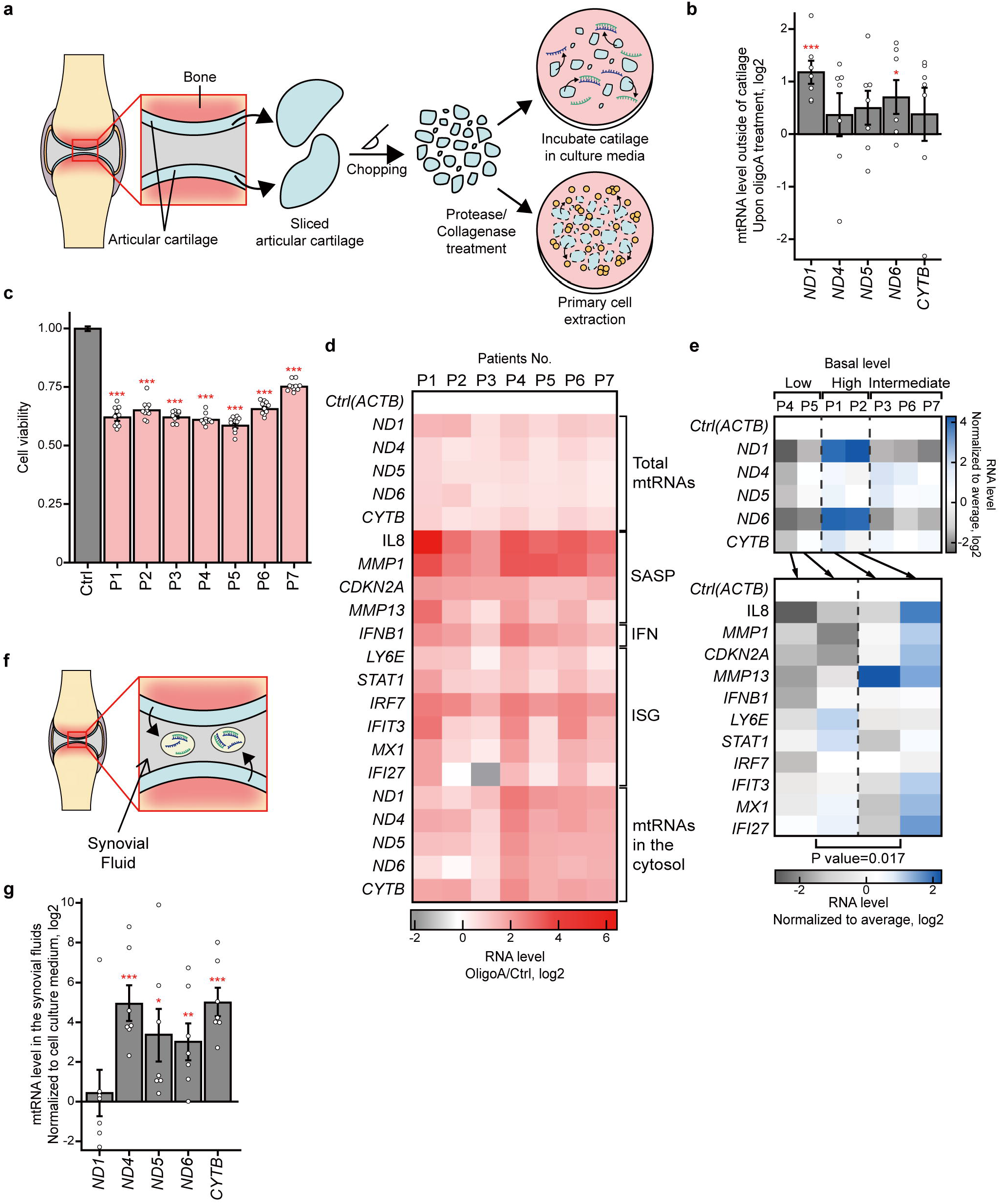
Analysis of cartilages, primary chondrocytes, and synovial fluids from OA patients. **a** A scheme of the experimental procedure for preparing cartilages and isolating primary chondrocytes from OA patients. **b** MRC inhibition in primary cartilages also results in extracellular export of mtRNAs. *n* = 7 and error bars denote s.e.m. **c** OligoA treatment in primary chondrocytes isolated from seven different OA patients resulted in significant cell death. *n* = 10 and error bars denote s.e.m. **d** Oligomycin A treatment in primary chondrocytes resulted in the elevation of total mtRNAs, induction of SASP factors, IFNs, ISGs, and cytosolic efflux of mtRNAs. *n* = 3 and error bars denote s.e.m. **e** Primary chondrocytes exhibit different basal levels of mtRNAs. Two patients with high basal mtRNA expression show higher expression of SASP factors, IFN, and ISGs compared to those of primary chondrocytes with low basal mtRNA expressions. *n* = 3 and error bars denote s.e.m. **f** A schematic representation of the synovial fluid. **g** Expression of mtRNAs from the synovial fluid normalized by basal mtRNA expression in media of SW1353 cells. *n* = 7 and error bars denote s.e.m.

We further processed cartilages by treating them with proteases and collagenases to isolate primary chondrocytes (Fig. 5a). We treated primary chondrocytes with oligoA and confirmed that inhibition of MRC leads to a significant cell death (Fig. 5c). Under this condition, we examined the expression and subcellular localization of mtRNAs as well as the induction of SASP factors and ISGs. We found that in primary chondrocytes, inhibition of MRC led to increased expression of total mtRNAs (Fig. 5d). Interestingly, samples with lower basal mtRNA levels resulted in a more pronounced induction of mtRNAs upon oligoA treatment. In addition, in all samples examined, oligoA treatment resulted in the significant cytosolic efflux of mtRNAs and induction of key SASP factors, including *IL8* and *MMP1* as well as *IFNβ1* and numerous ISGs (Fig. 5d). Furthermore, we closely examined the basal expression of mtRNAs in primary chondrocytes and found that cytosolic mtRNA levels before the oligoA treatment were correlated with the basal expressions of SASP factors and ISGs (Fig. 5e). Of note, the analyzed primary chondrocytes were extracted from the articular cartilages of OA patients. Therefore, our data indicate that cytosolic mtRNA expression may be used as an indicator to identify patients with activated dsRNA antiviral signaling and subsequent proinflammatory cytokine induction.

Lastly, we analyzed mtRNA expression in the synovial fluid of the patients. We hypothesized that if damaged chondrocytes releases mtRNAs to the extracellular space, then the level of mtRNAs in the synovial fluid should be elevated (Fig. 5f). We extracted RNAs from synovial fluids of seven different patients and performed RT-qPCR to examine mtRNA expression. We were able to detect mtRNAs clearly, but could not test whether or not the level was elevated in the patients because we could not acquire synovial fluids from healthy donors. Instead, we utilized the basal mtRNA expression from the media of SW1353 cells. To compare the two conditions, we normalized mtRNA expressions with that of *GAPDH*. We found that compared to the mtRNAs in cell culture media, the levels of mtRNAs normalized to that of *GAPDH* in the synovial fluids of the OA patients were significantly elevated (Fig. 5g). Overall, our analyses of OA patient samples underline the importance of mtRNAs in the development of OA.

### Autophagy alleviates OA phenotypes partly by removing cytosolic mtRNAs

Lastly, we investigated the therapeutic potential of targeting mtRNAs in protecting chondrocytes from mitochondrial dysfunction and in preventing subsequent development of OA. In particular, we asked whether the protective effect of autophagy, suggested by previous studies, is partly mediated through the removal of cytosolic mt-dsRNAs. Autophagy is a self digestion system that is essential to maintain cellular homeostasis in response to various stimuli like nutrient deprivation, infection, and programmed cell death. Recently, decreased autophagy activation due to aging has been shown to contribute to OA pathogenesis^50^. Indeed, treating chondrocytes with autophagy inducers such as rapamycin and torin-1 protected cells from mitochondrial dysfunction^51,52^. Yet, the mechanism behind the protective effect of autophagy during mitochondrial dysfunction is not fully understood. In light of our results and considering the working mechanism of autophagy, that it removes damaged macromolecules and organelles, we examined whether autophagy can remove cytosolic mt-dsRNAs to protect chondrocytes from mitochondrial damage.

To regulate autophagy induction, we triggered or blocked autophagy by treating cells with torin-1 or chloroquine (CQ), respectively. Autophagy activation was confirmed based on the decreased expression of microtubule-associated protein 1 light chain 3B-I (LC3B-I) and LC3B-II proteins (Supplementary Fig. 5a, b). In this study, we focused on the induction of autophagy using torin-1. Consistent with previous studies, activating autophagy rescued cell death from oligoA treatment (Supplementary Fig. 5c). More importantly, we found that when autophagy was activated, the levels of cytosolic mtRNAs in oligoA treated SW1353 cells were significantly decreased (Fig. 6a). Moreover, decreased cytosolic mtRNAs was translated into decreased expressions of *IL8* and *MMP1* as well as *IFNβ1* and several ISGs (Fig. 6b).

**Fig. 6.**
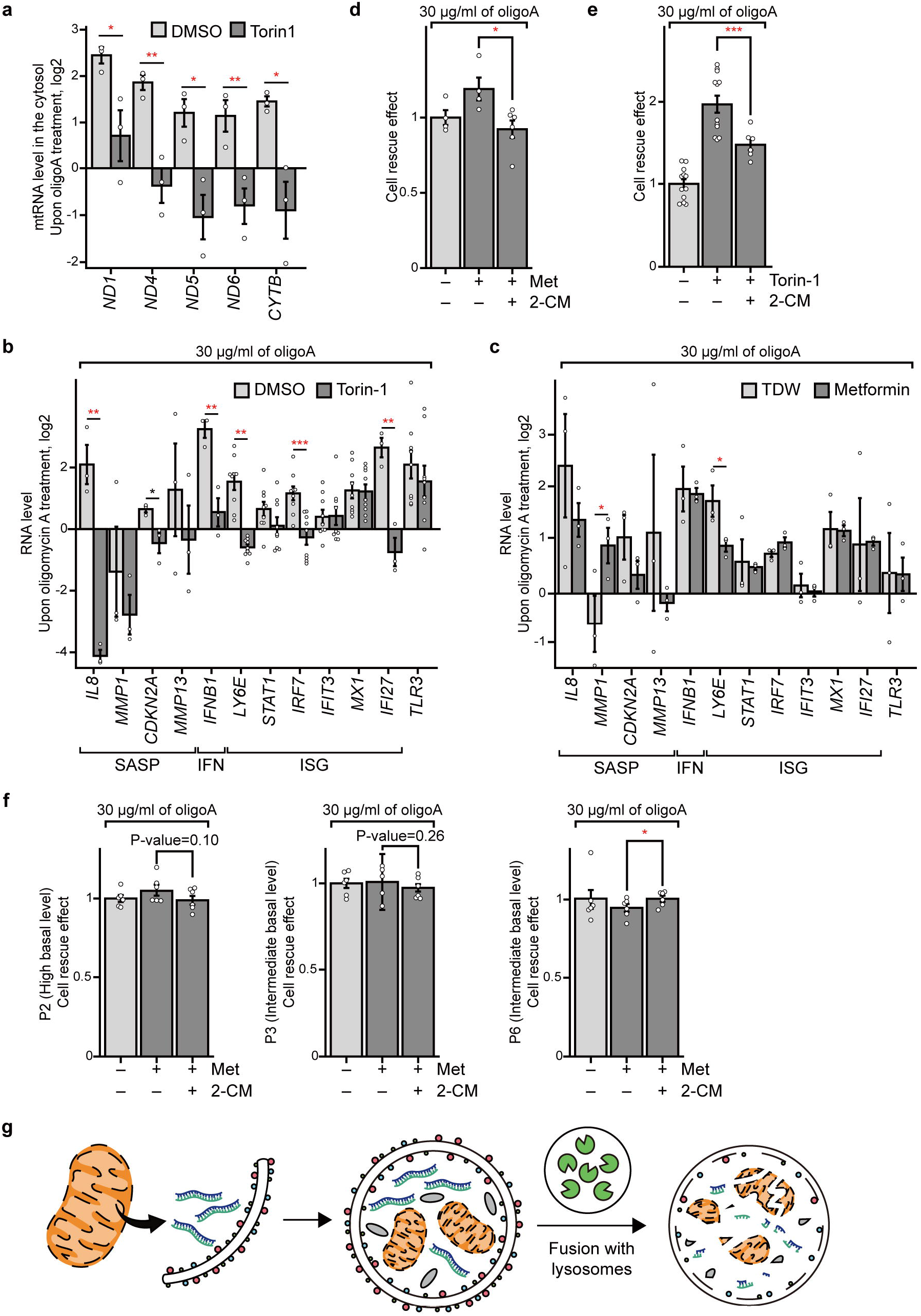
Autophagy alleviates OA partly by removing mtRNAs in the cytosol. **a, b** Torin-1 pretreatment decreases the expression of **a** free cytosolic mtRNAs and **b** several SASP and ISGs. *n* = 3 for cytosolic mtRNAs and SASP factors, *n* = 9 for ISGs; error bars denote s.e.m. **c** Inducing autophagy using metformin also rescues SASP factors and ISG expressions induced by oligoA treatment (*n* = 3). **d, e** The rescue effect of **d** metformin and **e** torin-1 pretreatment is decreased in cells with decreased mtRNA expression. *n* = 4 for metformin, *n* = 12 for torin-1 and error bars denote s.e.m. **f** The rescue effect of metformin pretreatment in primary chondrocytes isolated from the OA patients (*n* = 6). **g** A model for the removal of cytosolic mtRNAs by autophagy.

As an alternative way of activating autophagy, we treated SW1353 cells with metformin, an anti-diabetic drug that triggers autophagy by activating the AMPK pathway. Similar to the case of torin-1, metformin pretreatment also partly rescued cell death from mitochondrial dysfunction (Supplementary Fig. 5d). Moreover, metformin treatment also resulted in decreased expression of *IL8*, *MMP13*, and other SASP factors as well as several ISGs (Fig. 6c). The magnitude of the change was smaller for the case of metformin compared to those of torin-1, but the general effects were the same. To further investigate the importance of mtRNAs in the protective effect of autophagy from mitochondrial dysfunction, we performed the same cell viability experiment, but this time, we also pretreated cells with 2-CM to decrease the expression of mtRNAs. We found that in mtRNA-deficient cells, the protective effect of metformin was abrogated (Fig. 6d). In the case of torin-1, rescue effect was still observed, but the degree of the rescue was significantly reduced (Fig. 6e). Additionally, we analyzed whether the effect of metformin in primary chondrocyte cells isolated from the OA patients correlates with the basal mtRNA expressions. We found that metformin pretreatment showed rescue effect in a patient with high basal mtRNA expression (Fig. 6f). However, two patients with intermediate mtRNA expressions did not respond to metformin pretreatment. Therefore, autophagy protects chondrocytes from mitochondrial dysfunction partly by removing cytosolic mtRNAs released from damaged mitochondria (Fig. 6g).

### mt-dsRNAs are key signaling molecules in the development of OA

Taken together, our study identifies mt-dsRNAs as key signaling molecules that are involved in a wide range of processes that lead to the development of primary OA. According to our model, the cytosolic efflux of mt-dsRNAs and subsequent activation of RNA sensors of the innate immune response proteins such as PKR play a crucial role during chondrocyte senescence and in response to mitochondrial dysfunction by regulating the expression of SASP factors (Fig. 7). Moreover, mt-dsRNAs can also be exported outside the cells, where they act as autocrine-type signaling molecules to activate TLR3. In this context, our data suggest mtRNAs as the unknown dsRNAs from damaged articular chondrocytes to promote cartilage degeneration as suggested previously (Fig. 7). Indeed, targeting mtRNAs protects chondrocytes from mitochondrial dysfunction, and inducing autophagy alleviates OA progression partly through the removal of mtRNAs (Fig. 7). Therefore, our study establishes mt-dsRNAs as a critical regulator of OA development and suggests mt-dsRNAs as a potential therapeutic target to overcome OA.

**Fig. 7.**
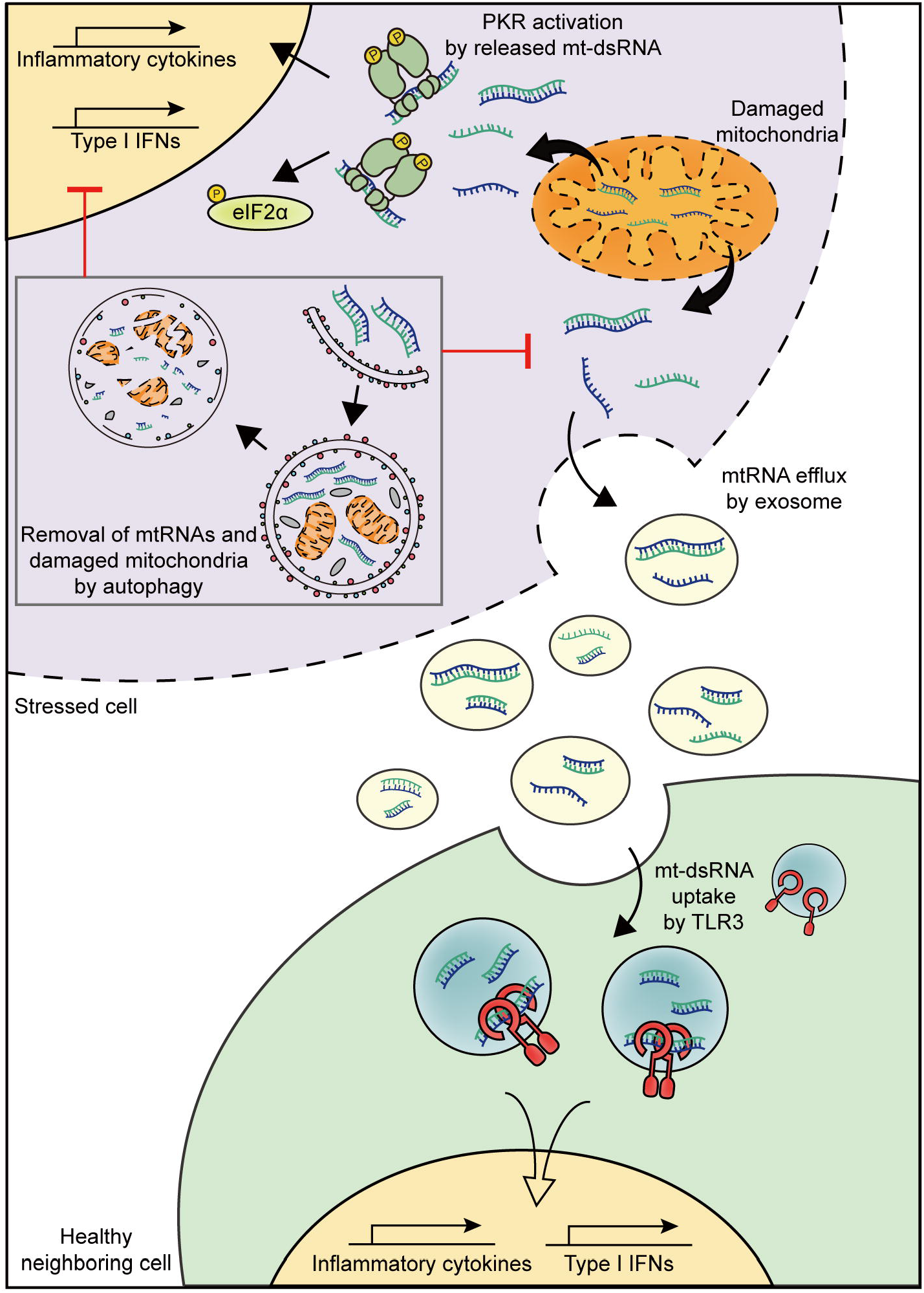
A model for the role of mtRNAs during OA pathogenesis. Mitochondrial dysfunction and chondrocyte senescence promote the cytosolic efflux of mtRNAs. The released mt-dsRNAs can activate the PKR signaling pathway, resulting in the stimulation of inflammatory cytokines and cell death (upper right). This can be prevented by the removal of cytosolic mtRNAs, which can occur through autophagy (top left). During mitochondrial dysfunction, mtRNAs can also be exported to the extracellular space where they act as antigens for TLR3 to promote cartilage degeneration (bottom).

## Discussion

A large part of the human genome consists of retrotransposons and repeat elements that are transcribed into RNAs but do not encode proteins. Recent evidence suggests that these non-coding elements can generate RNAs with long double-stranded secondary structures that are capable of activating innate immune response proteins that recognize viral dsRNAs ^35^. In addition, bidirectional transcription of the mitochondrial genome also results in the transcription of pairs of complementary RNAs that bind with each other and create long dsRNAs that can activate antiviral machinery^32,39^. However, the biological significance of mt-dsRNAs, particularly in the context of human disease, remains unknown. In this study, we establish that mt-dsRNAs as key intra- and intercellular signaling cues responsible for the activation of PKR and TLR3 to promote chondrocyte degeneration during the development of primary OA.

By investigating mt-dsRNA-mediated activation of PKR and TLR3, our study provides insights into understanding the pathogenesis and potential treatment of OA. Numerous studies reported increased levels of pPKR in damaged cartilages of OA patients^14,15,17^. Moreover, PKR activation was responsible for inflammation and MMP-13 secretion in degenerated articular chondrocytes^21^. The secretion of dsRNAs capable of activating TLR3 further supports the notion that PRRs are critical regulators during OA development. Yet, the activating cue of PKR and TLR3 remains unidentified. Our study suggests that during mitochondrial dysfunction and under senescence eliciting stress conditions, cytosolic efflux of mtRNAs is facilitated, and these cytosolic mtRNAs interact with PKR to activate the kinase. More importantly, decreased expression of mtRNAs phenocopied the effects of PKR knockdown, indicating that mt-dsRNAs are a primary activating source of PKR. Our study also provides a valuable insight into the role of mt-dsRNAs in promoting cartilage degeneration from damaged articular chondrocytes. In this context, under mitochondrial stress, mt-dsRNAs are exported to the extracellular space where they are recognized by TLR3 receptors in the neighboring cells to induce an inflammatory response. A similar phenomenon was observed when liver cells were under acute alcoholic stress^40^, indicating that mt-dsRNAs acting as intercellular signaling molecules might be a general response to various mitochondrial stressors.

Recently, many studies showed that activating autophagy can protect articular chondrocytes from mitochondrial stress and can alleviate OA symptoms^50,51,53^. Moreover, decreased activity of autophagy during aging has been suggested as a potential cause leading to the onset of OA. Yet, the underlying mechanism of the observed protective effect of autophagy was not fully understood. We show that autophagy rescues chondrocyte death from mitochondrial dysfunction partly by removing cytosolic mt-dsRNAs. Consistent with our idea, the effect of autophagy was reduced in mtRNA-deficient cells. Currently, it is unclear how autophagosomes may recognize cytosolic mtRNAs. One possibility is the random engulfment and degradation of cytosolic mtRNAs accidentally protected chondrocytes. In the future, it would be interesting to develop methods to induce autophagy to selectively remove cytosolic mt-dsRNAs as a potential treatment strategy of OA.

In addition to mtRNAs, retrotransposons and repeat elements can be transcribed into dsRNAs that can activate PKR and MDA5. Previous studies have shown that Alu elements occupy a substantial fraction of dsRNAome in human cells^32,54^. Indeed, many dsRNA-binding proteins such as adenosine deaminase acting on RNA (ADAR), STAUFEN, DHX9, PKR, and MDA5 can all recognize Alu dsRNAs^32,55–59^. Moreover, overactivation of MDA5 by Alus is associated with an autoimmune disease Aicardi-Goutières syndrome^59^, and increased LINE-1 expression is shown to contribute to OA development^37^. Collectively, mounting evidence suggests that retrotransposons may also act as cellular dsRNAs that activate PKR and TLR3 to promote cartilage degeneration. In the future, it will be interesting to uncover the possible function of retrotransposons and mt-dsRNAs that act as endogenous dsRNAs in the development of OA. Considering that the expression of retrotransposons is usually silenced in most cells, targeting these cellular dsRNAs may provide a new therapeutic direction in fighting OA. In sum, our study highlights the importance of mt-dsRNAs during OA development and presents selective removal of cellular dsRNAs as a possible treatment of degenerative diseases such as OA.

## Methods

### Cell culture and chemical treatment

SW1353 (ATCC) cells were grown in Dulbecco’s modified eagle medium (DMEM; Welgene) supplemented with 9.1% (v/v) fetal bovine serum (FBS). For CHON-001 (ATCC), 0.1 mg/mL of G418 (Invivogen) was also added to the media. Both cell lines were grown at 37°C under 5% CO_2_. To induce mitochondrial stress, 30 μg/mL of oligoA (Sigma Aldrich) was treated for 24 h. To trigger or block autophagy, cells were treated with 50 nM torin-1 (LC laboratory) or 50 μM chloroquine (Sigma Aldrich) for 4 h before oligomycin A treatment, respectively. 200 μM of metformin (Sigma Aldrich) was treated for 48 h to provoke autophagy. To induce the senescence, cells were incubated with 100 nM of doxorubicin (LC laboratory) for 3 days or 500 μM of hydrogen peroxide (Sigma Aldrich) for 4 days. To downregulate mtRNA expression, 50 μM of 2-CM (Santa Cruz Biotechnology) was treated for 48 h. To knockdown PKR expression, a mixture of four different siRNAs was transfected using Lipofectamine 3000 (Thermo Fisher Scientific) following the manufacturer’s instructions. Sequences of the siRNAs used in this study are provided in **Table S1**.

### Cartilage and primary cell culture of OA patients

Human cartilages were obtained from seven OA patients with a mean age of 74.3 ± 5.01 years. Synovium, lipid, and bone were removed to prepare cartilages. To culture, cartilages were first chopped using surgical blades in phosphate-buffered saline (PBS) supplemented with 1% (v/v) Penicillin-Streptomycin (Welgene). Cartilages were then washed with PBS, transferred into a 24-well plate (200 mg per well), and cultured in DMEM supplemented with 9.1% (v/v) FBS (Gibco) and 1% Antibiotic-Antimycotic (Thermo Fisher Scientific). Cartilages were incubated for 3 days before further analysis. To isolate primary chondrocytes, chopped cartilage was incubated in DMEM supplemented with 0.2% (w/v) protease (Sigma Aldrich) for 1 h in a 37°C and 5% CO_2_ incubator. Cartilages were then washed once with PBS and treated with 0.3% (w/v) collagenase (Sigma Aldrich) for 3 h to digest the collagen. Primary cells were filtered using a 70 μm cell strainer (Falcon) and washed once with PBS. Primary cells were incubated for 7 days in DMEM supplemented with 9.1% FBS and 1% Antibiotic-Antimycotic. This protocol was approved by the Seoul National University Bundang Hospital Institutional Review Board (IRB approval number: B-0607/035-018).

### Quantitative real-time PCR

Cytosolic RNAs were isolated from cell lysates using subcellular protein fractionation kit (Thermo Fisher Scientific) following the manufacturer’s protocol. TRIzol LS (Ambion) was added at a 3:1 ratio to the cytosolic fraction to extract the RNAs. To analyze RNAs released from the cells, media was collected by taking the supernatant after centrifugation at 10,000 × *g* for 30 sec. TRIzol LS was added into the supernatant to extract the nucleic acids. After precipitation, DNase I (TaKaRa) was treated to remove DNA, and purified RNA was reverse transcribed using RevertAid reverse transcriptase (Thermo Fisher Scientific). For RNAs extracted from cartilages or primary cells, SuperScript IV reverse transcriptase (Invitrogen) was used to synthesize cDNA. For strand-specific qRT-PCR, reverse transcription primers containing CMV promoter sequences were designed to target the specific gene. Primers used in the study are provided in **Tables S2-S4**.

### Immunocytochemistry and RNA fluorescent *in situ* hybridization

Cells were grown in a 1% (w/v) gelatin-coated confocal dish (SPL) for at least overnight. Cells were washed once with PBS and fixed using 4% (w/v) paraformaldehyde (Sigma Aldrich) for 10 min at room temperature. The fixed cells were permeabilized in 0.1% (v/v) Triton X-100 (Promega) diluted in PBS for 10 min and blocked in 1% (w/v) bovine serum albumin (BSA; RMBIO) for 1 h. Cells were incubated with primary antibodies diluted in 1% BSA for 1 h at room temperature. Cells were then washed 4 times with 0.1% (v/v) Tween-20 in PBS (PBST) and incubated with Alexa fluor fluorophore-labeled secondary antibodies (Thermo Fisher Scientific). DAPI was added in the secondary antibody solution to stain the nuclei. The primary antibodies used in this study are as follows: pPKR and PKR antibodies were purchased from Santa Cruz Biotechnology, and peIF2α, eIF2α, and LC3B antibodies were purchased from Cell Signaling Technology.

The mitochondrial membrane potential was analyzed using JC-1 dye (Abcam). Cells were washed with PBS once and incubated in pre-warmed media containing 5 μg/mL of JC-1 dye for 30 min in a 37°C and 5% CO_2_; incubator. Cells were then washed 3 times with PBS and imaged using Leica DMi8 fluorescence microscope with a 20x objective (NA = 0.40).

RNAScope RNA fluorescent *in situ* hybridization (ACD) was used to visualize mtRNA expressions. RNAScope probes for ND4 and ND5 mtRNAs were purchased from ACD, Inc. The pretreatment, hybridization, and signal amplification were done following the manufacturer’s instructions. To visualize mitochondria, cells were incubated with 1.5 μM of MitoGreen (PromoCell) for 15 min at room temperature. Zeiss LSM 780 confocal microscope using a 63x objective (NA = 1.40) was used for imaging.

### β-Galactosidase staining

The senescent status of the cells was analyzed using the β-Galactosidase staining kit (Cell Signaling Technology) following the manufacturer’s protocol. The stained cells were imaged using a 10x objective of Leica DMi1 microscope with NA = 0.22.

### PKR fCLIP

To prepare PKR antibody-conjugated beads, protein A beads (Thermo Fisher Scientific) was incubated with PKR antibody (Cell Signaling Technology) in the fCLIP lysis buffer (20 mM Tris-HCl, pH 7.5, 15 mM NaCl, 10 mM EDTA, 0.5% NP-40, 0.1% Triton X-100, 0.1% SDS, and 0.1% sodium deoxycholate) for 3 h at 4°C after adjusting NaCl concentration to 150 mM. Harvested cells were fixed with 0.1% (w/v) filtered paraformaldehyde (Sigma Aldrich) for 10 min and immediately quenched by adjusting glycine (Bio-basic) concentration to 250 mM. The crosslinked cells were lysed using the fCLIP lysis buffer for 10 min on ice and then sonicated. The NaCl concentration of the lysate was adjusted to 150 mM, and cell debris was separated by centrifugation. The lysate was added to the PKR antibody-conjugated beads and incubated for 3 h at 4°C. The bead was washed 4 times with the fCLIP wash buffer (20 mM Tris-HCl, pH 7.5, 150 mM NaCl, 10 mM EDTA, 0.1% NP-40, 0.1% SDS, 0.1% Triton X-100, and 0.1% sodium deoxycholate) and PKR-dsRNA complex was eluted from the beads by incubating in the elution buffer (200 mM Tris-HCl pH 7.4, 100 mM NaCl, 20 mM EDTA, 2% SDS, and 7 M Urea) for 3 h at 25°C. The eluate was treated with 20 mg/ml proteinase K (Sigma Aldrich) for overnight at 65°C. RNA was purified using acid phenol:Chloroform pH 4.5 (Thermo Fisher Scientific).

### Western blotting

Cell lysates were prepared by incubating cells in the lysis buffer (50 mM Tris-HCl pH 8.0, 100 mM KCl, 0.5% NP-40, 10% Glycerol, and 1 mM DTT) followed by sonication. 30~50 μg of protein samples were separated on a 10% SDS-PAGE gel and transferred to a PVDF membrane (Merck) using an Amersham semi-dry transfer system. The primary antibodies used in this study were as follows: pPKR (Abcam), Lamin A/C (Santa Cruz Biotechnology), PKR, eIF2α, peIF2α, LC3B, β-tubulin, JNK, AIF, RAB5, and Histone H3 (Cell signaling Technology).

### Sulforhodamine B (SRB) assay

Cells grown in a 24-well plate were fixed with pre-chilled 10% (w/v) trichloroacetic acid (Sigma Aldrich) solution in distilled water for 1 h at 4°C. Cells were washed 4 times with cold PBS and air-dried. Cells were stained with 0.4% (w/v) sulforhodamine B (Santa Cruz Biotechnology) solution in 1% acetic acid for 30 min at room temperature. The plate was rinsed with 1% acetic acid and dried while protecting from light. The dye was eluted using 10 mM Tris solution (pH 10.5) and transferred to a 96-well plate. The absorbance at 510 nm was measured using Varioskan™ LUX multimode microplate reader (Thermo Fisher Scientific).

### Statistical analysis

Quantitative RT-PCR data and SRB assay results were analyzed using the one-tailed Student’s t-test. All data were biologically replicated for at least 3 times. The error bars indicate the standard error of the mean. p-values ≤ 0.05 were regarded as statistically significant. * denotes p-values ≤ 0.05, ** is p-values ≤ 0.01, and *** is p-values ≤ 0.001.

## Reporting summary

Further information on research design is available in the Nature Research Reporting Summary linked to this article.

## Data availability

The datasets used and/or analyzed during the current study are available from the corresponding authors on reasonable request.

## Code availability

This study did not use custom code or mathematical algorithms.

## Acknowledgments

We thank all members of the Lee and Kim laboratory for helpful discussion and comments on the paper.

## Author contributions

S. Kim, Y. Lee, and Y. Kim designed the research. S. Kim and K. Lee performed the most experiments with help from Y. Choi. All authors analyzed the data. S. Kim, K. Lee, Y. Lee, and Y. Kim wrote the paper.

## References

1. Ehrenfeld, E. & Hunt, T. Double-stranded poliovirus RNA inhibits initiation of protein synthesis by reticulocyte lysates. Proc Natl Acad Sci U S A 68, 1075–8 (1971).

2. Levin, D.H., Petryshyn, R. & London, I.M. Characterization of double-stranded-RNA-activated kinase that phosphorylates alpha subunit of eukaryotic initiation factor 2 (eIF-2 alpha) in reticulocyte lysates. Proc Natl Acad Sci U S A 77, 832–6 (1980).

3. Dey, M. et al. Mechanistic link between PKR dimerization, autophosphorylation, and eIF2alpha substrate recognition. Cell 122, 901–13 (2005).

4. Patel, R.C., Stanton, P., McMillan, N.M., Williams, B.R. & Sen, G.C. The interferon-inducible double-stranded RNA-activated protein kinase self-associates in vitro and in vivo. Proc Natl Acad Sci U S A 92, 8283–7 (1995).

5. Lemaire, P.A., Anderson, E., Lary, J. & Cole, J.L. Mechanism of PKR Activation by dsRNA. J Mol Biol 381, 351–60 (2008).

6. Meurs, E.F. et al. Constitutive expression of human double-stranded RNA-activated p68 kinase in murine cells mediates phosphorylation of eukaryotic initiation factor 2 and partial resistance to encephalomyocarditis virus growth. J Virol 66, 5805–14 (1992).

7. Takada, Y., Ichikawa, H., Pataer, A., Swisher, S. & Aggarwal, B.B. Genetic deletion of PKR abrogates TNF-induced activation of IkappaBalpha kinase, JNK, Akt and cell proliferation but potentiates p44/p42 MAPK and p38 MAPK activation. Oncogene 26, 1201–12 (2007).

8. Garcia, M.A. et al. Impact of protein kinase PKR in cell biology: from antiviral to antiproliferative action. Microbiol Mol Biol Rev 70, 1032–60 (2006).

9. Kim, Y. et al. PKR is activated by cellular dsRNAs during mitosis and acts as a mitotic regulator. Genes Dev 28, 1310–22 (2014).

10. Kim, Y. et al. Deletion of human tarbp2 reveals cellular microRNA targets and cell-cycle function of TRBP. Cell Rep 9, 1061–74 (2014).

11. Zhu, P.J. et al. Suppression of PKR promotes network excitability and enhanced cognition by interferon-gamma-mediated disinhibition. Cell 147, 1384–96 (2011).

12. Dumurgier, J. et al. Cerebrospinal fluid PKR level predicts cognitive decline in Alzheimer’s disease. PLoS One 8, e53587 (2013).

13. Bando, Y. et al. Double-strand RNA dependent protein kinase (PKR) is involved in the extrastriatal degeneration in Parkinson’s disease and Huntington’s disease. Neurochem Int 46, 11–8 (2005).

14. Gilbert, S.J., Duance, V.C. & Mason, D.J. Does protein kinase R mediate TNF-alpha- and ceramide-induced increases in expression and activation of matrix metalloproteinases in articular cartilage by a novel mechanism? Arthritis Res Ther 6, R46–R55 (2004).

15. Gilbert, S.J. et al. Deletion of P58(IPK), the Cellular Inhibitor of the Protein Kinases PKR and PERK, Causes Bone Changes and Joint Degeneration in Mice. Front Endocrinol (Lausanne) 5, 174 (2014).

16. Ohno, M. Roles of eIF2alpha kinases in the pathogenesis of Alzheimer’s disease. Front Mol Neurosci 7, 22 (2014).

17. Tam, C.L., Hofbauer, M. & Towle, C.A. Requirement for protein kinase R in interleukin-1alpha-stimulated effects in cartilage. Biochem Pharmacol 74, 1636–41 (2007).

18. Oliveria, S.A., Felson, D.T., Reed, J.I., Cirillo, P.A. & Walker, A.M. Incidence of symptomatic hand, hip, and knee osteoarthritis among patients in a health maintenance organization. Arthritis Rheum 38, 1134–41 (1995).

19. Burrage, P.S., Mix, K.S. & Brinckerhoff, C.E. Matrix metalloproteinases: role in arthritis. Front Biosci 11, 529–43 (2006).

20. Yang, C.Y., Chanalaris, A. & Troeberg, L. ADAMTS and ADAM metalloproteinases in osteoarthritis - looking beyond the ‘usual suspects’. Osteoarthritis Cartilage 25, 1000–1009 (2017).

21. Ma, C.H. et al. PKR activation causes inflammation and MMP-13 secretion in human degenerated articular chondrocytes. Redox Biol 14, 72–81 (2018).

22. Li, C. et al. Double-stranded RNA released from damaged articular chondrocytes promotes cartilage degeneration via Toll-like receptor 3-interleukin-33 pathway. Cell Death Dis 8, e3165 (2017).

23. Guo, Z., Li, Y. & Ding, S.W. Small RNA-based antimicrobial immunity. Nat Rev Immunol 19, 31–44 (2019).

24. Weber, F., Wagner, V., Rasmussen, S.B., Hartmann, R. & Paludan, S.R. Double stranded RNA is produced by positive-strand RNA viruses and DNA viruses but not in detectable amounts by negative-strand RNA viruses. J Virol 80, 5059–64 (2006).

25. Tatematsu, M., Funami, K., Seya, T. & Matsumoto, M. Extracellular RNA Sensing by Pattern Recognition Receptors. J Innate Immun 10, 398–406 (2018).

26. Chen, L.L., DeCerbo, J.N. & Carmichael, G.G. Alu element-mediated gene silencing. EMBO J 27, 1694–705 (2008).

27. Athanasiadis, A., Rich, A. & Maas, S. Widespread A-to-I RNA editing of Alu-containing mRNAs in the human transcriptome. PLoS Biol 2, e391 (2004).

28. Lander, E.S. et al. Initial sequencing and analysis of the human genome. Nature 409, 860–921 (2001).

29. Deininger, P.L. & Batzer, M.A. Mammalian retroelements. Genome Res 12, 1455–65 (2002).

30. Karijolich, J., Zhao, Y., Alla, R. & Glaunsinger, B. Genome-wide mapping of infection-induced SINE RNAs reveals a role in selective mRNA export. Nucleic Acids Res 45, 6194–6208 (2017).

31. Elbarbary, R.A., Lucas, B.A. & Maquat, L.E. Retrotransposons as regulators of gene expression. Science 351, aac7247 (2016).

32. Kim, Y. et al. PKR Senses Nuclear and Mitochondrial Signals by Interacting with Endogenous Double-Stranded RNAs. Mol Cell 71, 1051–1063 e6 (2018).

33. Chiappinelli, K.B. et al. Inhibiting DNA Methylation Causes an Interferon Response in Cancer via dsRNA Including Endogenous Retroviruses. Cell 162, 974–86 (2015).

34. Pham, A.M. et al. PKR Transduces MDA5-Dependent Signals for Type I IFN Induction. PLoS Pathog 12, e1005489 (2016).

35. Kim, S., Ku, Y., Ku, J. & Kim, Y. Evidence of Aberrant Immune Response by Endogenous Double-Stranded RNAs: Attack from Within. Bioessays 41, e1900023 (2019).

36. Kaneko, H. et al. DICER1 deficit induces Alu RNA toxicity in age-related macular degeneration. Nature 471, 325–30 (2011).

37. Teerawattanapong, N., Udomsinprasert, W., Ngarmukos, S., Tanavalee, A. & Honsawek, S. Blood leukocyte LINE-1 hypomethylation and oxidative stress in knee osteoarthritis. Heliyon 5, e01774 (2019).

38. Barshad, G., Marom, S., Cohen, T. & Mishmar, D. Mitochondrial DNA Transcription and Its Regulation: An Evolutionary Perspective. Trends Genet 34, 682–692 (2018).

39. Dhir, A. et al. Mitochondrial double-stranded RNA triggers antiviral signalling in humans. Nature 560, 238–242 (2018).

40. Lee, J.H. et al. Mitochondrial double-stranded RNA in exosome promotes interleukin-17 production through toll-like receptor 3 in alcoholic liver injury. Hepatology (2019).

41. Liu, J.T. et al. Mitochondrial function is altered in articular chondrocytes of an endemic osteoarthritis, Kashin-Beck disease. Osteoarthritis Cartilage 18, 1218–26 (2010).

42. Ruiz-Romero, C. et al. Mitochondrial dysregulation of osteoarthritic human articular chondrocytes analyzed by proteomics: a decrease in mitochondrial superoxide dismutase points to a redox imbalance. Mol Cell Proteomics 8, 172–89 (2009).

43. Vaamonde-Garcia, C. et al. Mitochondrial dysfunction increases inflammatory responsiveness to cytokines in normal human chondrocytes. Arthritis Rheum 64, 2927–36 (2012).

44. Gavriilidis, C., Miwa, S., von Zglinicki, T., Taylor, R.W. & Young, D.A. Mitochondrial dysfunction in osteoarthritis is associated with down-regulation of superoxide dismutase 2. Arthritis Rheum 65, 378–87 (2013).

45. Vaamonde-Garcia, C. et al. The mitochondrial inhibitor oligomycin induces an inflammatory response in the rat knee joint. BMC Musculoskelet Disord 18, 254 (2017).

46. Loeser, R.F., Collins, J.A. & Diekman, B.O. Ageing and the pathogenesis of osteoarthritis. Nat Rev Rheumatol 12, 412–20 (2016).

47. Childs, B.G., Durik, M., Baker, D.J. & van Deursen, J.M. Cellular senescence in aging and age-related disease: from mechanisms to therapy. Nat Med 21, 1424–35 (2015).

48. Kang, D. et al. Stress-activated miR-204 governs senescent phenotypes of chondrocytes to promote osteoarthritis development. Sci Transl Med 11(2019).

49. Kunkeaw, N. et al. Mechanism mediated by a non-coding RNA, nc886, in the cytotoxicity of a DNA-reactive compound. Proc Natl Acad Sci U S A 116, 8289–8294 (2019).

50. Carames, B., Taniguchi, N., Otsuki, S., Blanco, F.J. & Lotz, M. Autophagy is a protective mechanism in normal cartilage, and its aging-related loss is linked with cell death and osteoarthritis. Arthritis Rheum 62, 791–801 (2010).

51. Cheng, N.T., Guo, A. & Meng, H. The protective role of autophagy in experimental osteoarthritis, and the therapeutic effects of Torin 1 on osteoarthritis by activating autophagy. BMC Musculoskelet Disord 17, 150 (2016).

52. Carames, B. et al. Autophagy activation by rapamycin reduces severity of experimental osteoarthritis. Ann Rheum Dis 71, 575–81 (2012).

53. Sasaki, H. et al. Autophagy modulates osteoarthritis-related gene expression in human chondrocytes. Arthritis Rheum 64, 1920–8 (2012).

54. Batzer, M.A. & Deininger, P.L. Alu repeats and human genomic diversity. Nat Rev Genet 3, 370–9 (2002).

55. Chung, H. et al. Human ADAR1 Prevents Endogenous RNA from Triggering Translational Shutdown. Cell 172, 811–824 e14 (2018).

56. Aktas, T. et al. DHX9 suppresses RNA processing defects originating from the Alu invasion of the human genome. Nature 544, 115–119 (2017).

57. Chu, W.M., Ballard, R., Carpick, B.W., Williams, B.R. & Schmid, C.W. Potential Alu function: regulation of the activity of double-stranded RNA-activated kinase PKR. Mol Cell Biol 18, 58–68 (1998).

58. Gong, C. & Maquat, L.E. lncRNAs transactivate STAU1-mediated mRNA decay by duplexing with 3’ UTRs via Alu elements. Nature 470, 284–8 (2011).

59. Ahmad, S. et al. Breaching Self-Tolerance to Alu Duplex RNA Underlies MDA5-Mediated Inflammation. Cell 172, 797–810 e13 (2018).

