## Supplementary figures for "Mitochondrial dsRNAs activate PKR and TLR3 to promote chondrocyte degeneration in osteoarthritis"

### SUPPLEMENTARY FIGURE LEGENDS

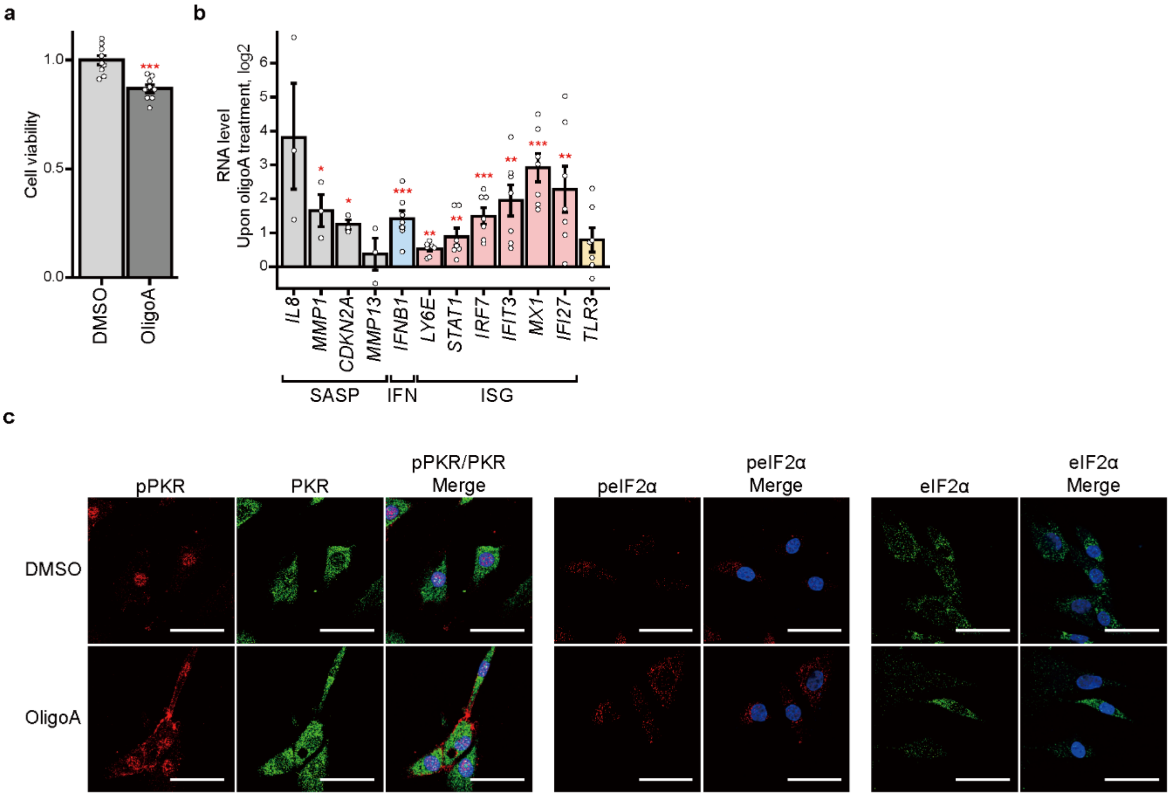


**Supplementary Fig. 1 MRC inhibition activates the PKR signaling in CHON-001 immortalized human chondrocytes. a** SRB assay analysis to show cell viability upon oligoA treatment. *n* = 9 and error bars denotes s.e.m. **b** mRNA expressions of SASP factors, IFN, and ISGs in response to oligoA treatment. *n* = 3 for SASP factors, *n* = 7 for IFN and ISGs and error bars denote s.e.m. **c** Immunocytochemistry staining of the PKR signaling pathway, following oligoA treatment. Scale bar, 50 μm.

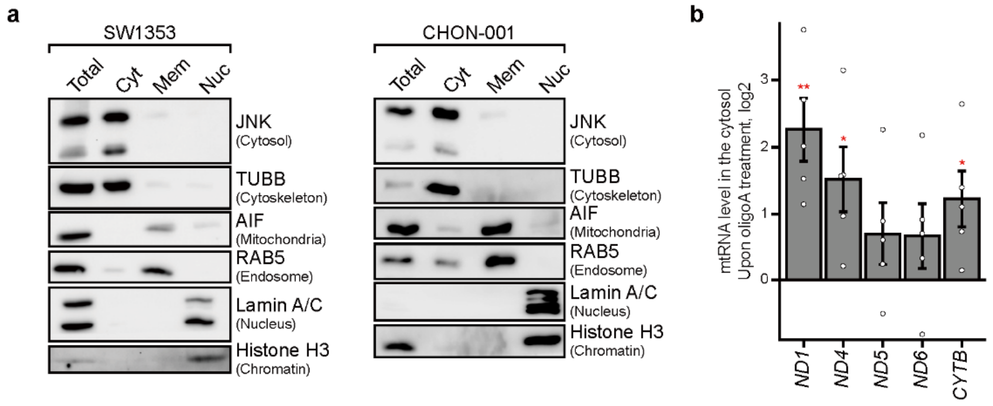


**Supplementary Fig. 2 Subcellular fractionation and cytosolic mtRNA expression. a** Verification of successful subcellular fractionation in SW1353 and CHON-001 using immunoblotting of key marker proteins. **b** Cytosolic mtRNA expression levels upon oligoA treatment in CHON-001 cells. *n* = 5 and error bars denote s.e.m.

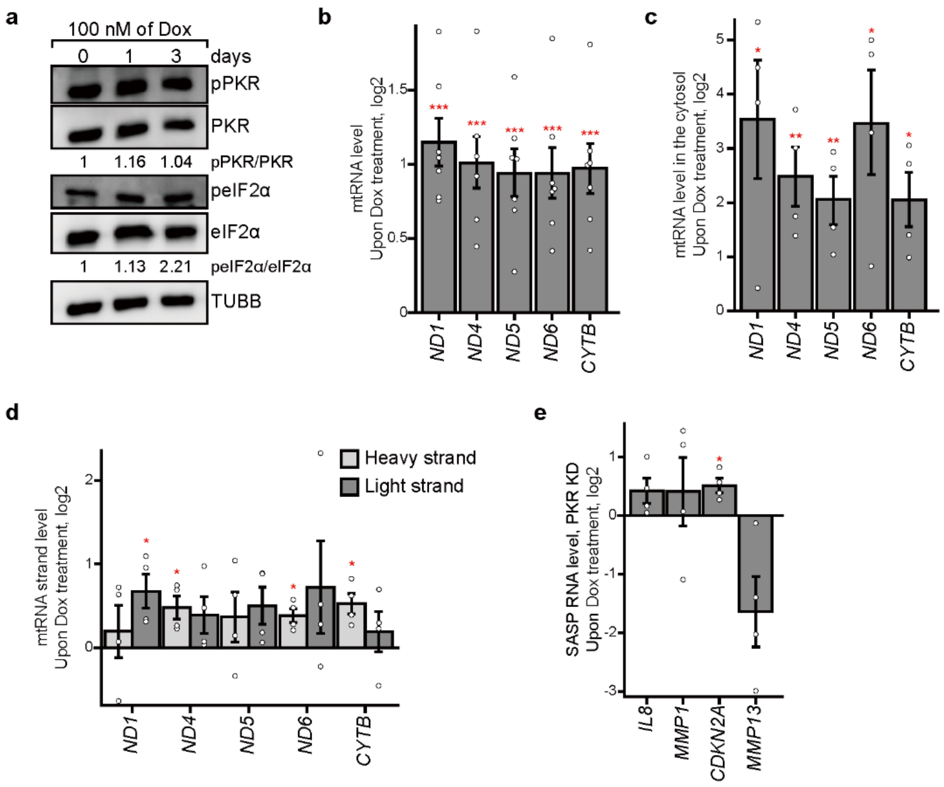


**Supplementary Fig. 3 Inducing senescence using doxorubicin activates PKR via mtRNAs.** **a** Activation of PKR and phosphorylation of eIF2α in response to Dox treatment over time. **b, c** **b** Total and **c** free cytosolic mtRNA expressions 3 days after Dox treatment. *n* = 7 for total, *n* = 4 for free cytosolic mtRNA expressions and error bars denote s.e.m. **d** Strand-specific analysis of mtRNAs indicate increased expression of both sense and antisense transcripts upon Dox treatment. *n* = 4 and error bars denote s.e.m. **e** Decreased expression of *MMP13* mRNA 3 days after Dox treatment in PKR-deficient cells (*n* = 4).

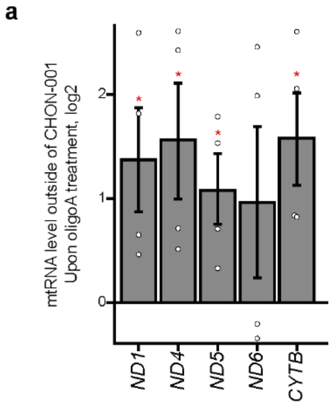


**Supplementary Fig. 4 mtRNA efflux to the extracellular space in CHON-001 cells. a** mtRNA efflux in CHON-001 to the extracellular space upon MRC inhibition. $n$ = 4 and error bars denote s.e.m.

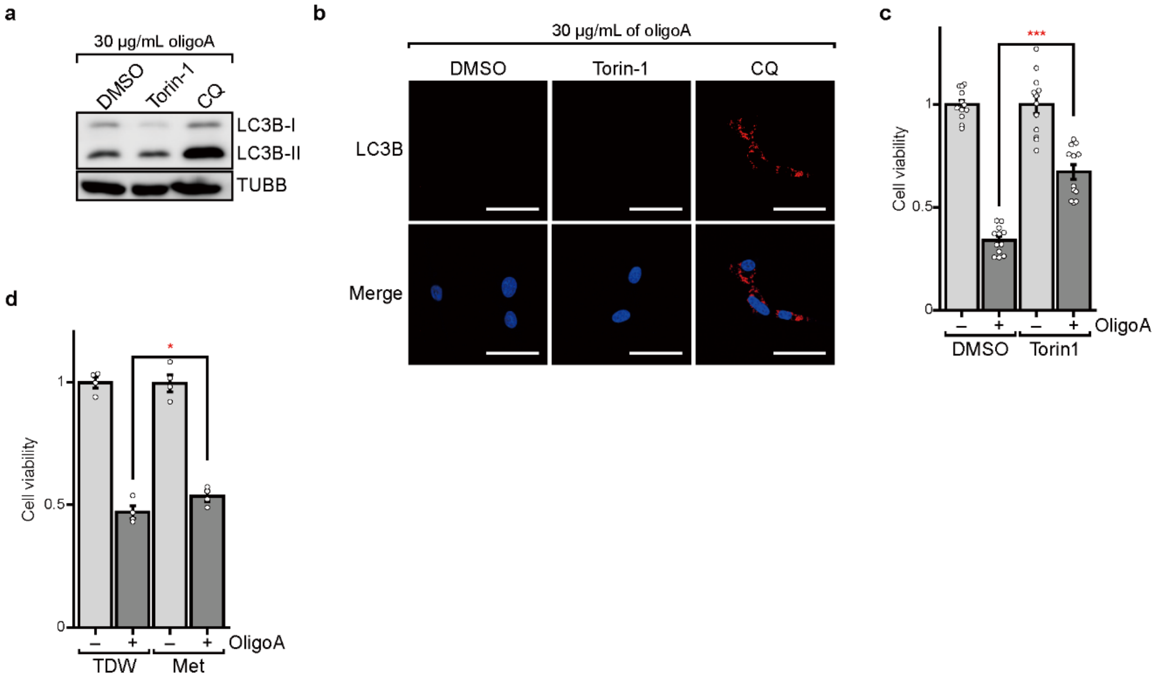


**Supplementary Fig. 5 Autophagy induction and chondrocyte rescue. a, b** Immunoblotting and immunocytochemistry analysis of LC3B expression upon inducing or inhibiting autophagy by using torin-1 or chloroquine (CQ), respectively. Scale bar, 50 µm. **c, d** Inducing autophagy through **c** torin-1 or **d** metformin pretreatment rescues cell death by MRC inhibition. *n* = 12 for torin-1 and *n* = 4 for metformin and error bars denote s.e.m.

**Table S1.** Target site sequences of siRNA

| Gene | Sequences (5′-3′) |
|---|---|
| siLuc | CUU ACG CUG AGU ACU UCG A |
| siPKR-1 | GCA GGG AGU AGU ACU UAA A |
| siPKR-2 | GCA UGG GCC AGA AGG AUU U |
| siPKR-3 | GCA GAU ACA UCA GAG AUA A |
| siTLR3 | ACU UUG CCU UGU AUC UAC U |

**Table S2.** Primer sequences for RT-qPCR

| Gene | Forward Primer (5′-3′) | Reverse Primer (5′-3′) |
|---|---|---|
| ND1 | TCAAACTCAAACTACGCCCTG | CGCAAATGGGCGGTAGGCGTG GTTGTGATAAGGGTGGAGAGG |
| ND4 | CTCACACTCATTCTCAACCCC | CGCAAATGGGCGGTAGGCGTGT GTTTGTCGTAGGCAGATGG |
| ND5 | CTAGGCCTTCTTACGAGCC | CGCAAATGGGCGGTAGGCGTGT TTGGGTTGAGGTGATGATG |
| ND6 | TGCTGTGGGTGAAAGAGTATG | CGCAAATGGGCGGTAGGCGTGC CCATAATCATACAAAGCCCC |
| CYTB | CAATTATACCCTAGCCAACCCC | CGCAAATGGGCGGTAGGCGTG GGATAGTAATAGGGCAAGGAC G |
| IL8 | ATGACTTCCAAGCTGGCCGTGG CT | TCTCAGCCCTCTTCAAAAACTT CTC |
| MMP1 | AGAAAGAAGACAAAGGCAAGT TGA | TTCCCAGTCACTTTCAGCCC |
| CDKN2A | ACTTCAGGGGTGCCACATTC | CGACCCTGTCCCTCAAATCC |
| MMP13 | GCCATTACCAGTCTCCGAGG | TACGGTTGGGAAGTTCTGGC |
| IFNB1 | TCTCCTGTTGTGCTTCTCCAC | GCCTCCCATTCAATTGCCAC |
| LY6E | CTCCAGGCAGGACGGCCATC | CGAGATTCCCAATGCCGGCACT |
| STAT1 | AACCTCGACAGTCTTGGCAC | CACTGAGACATCCTGCCACC |
| IRF7 | CTGTGGACACCTGTGACACC | TGCCCTCTCAGGAGCCAA |
| IFIT3 | GAAGGAACTGGGCCGCCTGCTA AG | GCCCTGGCCCATTTCCTCACTA CC |
| MX1 | TTCTGGGTCGGAGGCTACAG | TGGATGGCGGCGTTCT |
| IFI27 | ATCAGCAGTGACCAGTGTGG | TGGCCACAACTCCTCCAATC |
| TLR3 | TAGCAGTCATCCAACAGAATCA T | AATCTTCTGAGTTGATTATGGG TAA |
| IRF3 | ACACATACTGGGCAGTGAGC | CTACAATGAAGGGCCCCAGG |
| TRAF3 | ACCGCGAGAACTCCTCTTTC | TCAGGGACAAAAACTGGCGT |

| | | |
|---|---|---|
| IκBα | CTCCGAGACTTTCGAGGAAATAC | GCCATTGTAGTTGGTAGCCTTCA |
| ACTB | CCTGTACGCCAACACAGTGC | ATACTCCTGCTTGCTGATCC |
| GAPDH | CTCCTCCACCTTTGACGCTG | TCCTCTTGTGCTCTTGCTGG |

**Table S3. P**rimer sequences for strand-specific reverse transcription

| **Primer** | **Sequences (5′-3′)** |
|---|---|
| ND1 Heavy | CGCAAATGGGCGGTAGGCGTGGTTGTGATAAGGGTGGAGAGG |
| ND1 Light | CGCAAATGGGCGGTAGGCGTGTCAAACTCAAACTACGCCCTG |
| ND4 Heavy | CGCAAATGGGCGGTAGGCGTGTGTTTGTCGTAGGCAGATGG |
| ND4 Light | CGCAAATGGGCGGTAGGCGTGCCTCACACTCATTCTCAACCC |
| ND5 Heavy | CGCAAATGGGCGGTAGGCGTGTTTGGGTTGAGGTGATGATG |
| ND5 Light | CGCAAATGGGCGGTAGGCGTGCATTGTCGCATCCACCTTTA |
| ND6 Heavy | CGCAAATGGGCGGTAGGCGTGGGTTGAGGTCTTGGTGAGTG |
| ND6 Light | CGCAAATGGGCGGTAGGCGTGCCCATAATCATACAAAGCCCC |
| CYTB Heavy | CGCAAATGGGCGGTAGGCGTGGGATAGTAATAGGGCAAGGACG |
| CYTB Light | CGCAAATGGGCGGTAGGCGTGCAATTATACCCTAGCCAACCCC |
| GAPDH | CGCAAATGGGCGGTAGGCGTGTGAGCGATGTGGCTCGGCT |
| ACTB | CGCAAATGGGCGGTAGGCGTGACA CAG AGTACTTGCGCTCAG |

**Table S4.** Primer sequences for strand-specific qPCR

| Gene | Forward Primer (5′-3′) | Reverse Primer (5′-3′) |
|---|---|---|
| ND1 Heavy | TCAAACTCAAACTACGCCCTG | CGCAAATGGGCGGTAGGCGTG |
| ND1 Light | GTTGTGATAAGGGTGGAGAGG | CGCAAATGGGCGGTAGGCGTG |
| ND4 Heavy | CTCACACTCATTCTCAACCCC | CGCAAATGGGCGGTAGGCGTG |
| ND4 Light | TGTTTGTCGTAGGCAGATGG | CGCAAATGGGCGGTAGGCGTG |
| ND5 Heavy | CTAGGCCTTCTTACGAGCC | CGCAAATGGGCGGTAGGCGTG |
| ND5 Light | TAGGGAGAGCTGGGTTGTTT | CGCAAATGGGCGGTAGGCGTG |
| ND6 Heavy | TCATACTCTTTCACCCACAGC | CGCAAATGGGCGGTAGGCGTG |
| ND6 Light | TGCTGTGGGTGAAAGAGTATG | CGCAAATGGGCGGTAGGCGTG |
| CYTB Heavy | CAATTATACCCTAGCCAACCCC | CGCAAATGGGCGGTAGGCGTG |
| CYTB Light | GGATAGTAATAGGGCAAGGACG | CGCAAATGGGCGGTAGGCGTG |
| GAPDH | CAACGACCACTTTGTCAAGC | CGCAAATGGGCGGTAGGCGTG |
| ACTB | ACACAGTGCTGTCTCGTGGTA | CGCAAATGGGCGGTAGGCGTG |
